# Sensory Reweighting: A Common Mechanism for Subjective Visual Vertical and Cybersickness Susceptibility

**DOI:** 10.1101/2022.11.18.517106

**Authors:** William Chung, Michael Barnett-Cowan

## Abstract

The malaise symptoms of cybersickness are thought to be related to the sensory conflict present in the exposure to virtual reality (VR) content. When there is a sensory mismatch in the process of sensory perception, the perceptual estimate has been shown to change based on a reweighting mechanism between the relative contributions of the individual sensory signals involved. In this study, the reweighting of vestibular and body signals was assessed before and after exposure to different typical VR experiences and sickness severity was measured to investigate the relationship between susceptibility to cybersickness and sensory reweighting. Participants reported whether a visually presented line was rotated clockwise or counterclockwise from vertical while laying on their side in a subjective visual vertical (SVV) task. Task performance was recorded prior to VR exposure and after a low and high intensity VR game. The results show that the SVV was significantly shifted away from the body representation of upright and towards the vestibular signal after exposure to the high intensity VR game. Cybersickness measured using the fast motion sickness (FMS) scale found that sickness severity ratings were higher in the high intensity compared to the low intensity experience. The change in SVV from baseline after each VR exposure modelled using a simple 3-parameter gaussian regression fit was found to explain 49.5% of the variance in the FMS ratings. These results highlight the aftereffects of VR for sensory perception and suggests a potential relationship between the susceptibility to cybersickness and sensory reweighting.

## Introduction

Virtual reality (VR) technology has grown immensely and become widely available for both consumer entertainment and for applications in research and therapy (Bohil et al., 2011). Despite the advancement in VR displays and systems, cybersickness still remains a concern and a barrier to using VR technology for most users. Cybersickness consists of a variety of unpleasant symptoms related to nausea, disorientation, eye strain and fatigue and these symptoms can manifest differently depending on the individual (LaViola Jr, 2000; Rebenitsch and Owen, 2016). This has made it challenging for researchers to accurately quantify and predict the occurrence of sickness. Recent work using physiological and neuroimaging techniques have provided better insights into the neural basis of cybersickness (Cohen et al., 2019; Rebenitsch and Owen, 2016; Toschi et al., 2017). However, more research is still needed to understand the mechanisms by which cybersickness is triggered along with any interactions with the virtual content that the user is experiencing. Researchers have been able to predict between 34 to 55% of the incidence of cybersickness using various modelling approaches between sickness ratings and physiological measures, sensory perception and behaviours, demographic information, as well as hardware and software factors (Chang et al., 2021; Dennison et al., 2016; Kim et al., 2005; Nooij et al., 2017; Rebenitsch and Owen, 2021; Weech et al., 2018b).

One of the prevalent explanations of cybersickness is the sensory mismatch theory (Oman, 1990; Reason and Brand; 1975; Reason, 1978), which describes how sickness is induced when there is a discrepancy in feedback from the sensory inputs signaling self-motion. This conflict is notably evident when immersed in most virtual reality experiences in which there is an abundance of visual motion, but minimal movement signalled by the vestibular and proprioceptive systems leading to discomfort and sickness (Cobb et al., 1999; Howarth and Costello, 1997; Keshavarz et al., 2015; Kim et al., 2020). While there have been arguments against the sensory conflict theory (Stoffregen and Riccio, 1991) regarding the inconsistencies in predicting the occurrence and severity of sickness and how sensory mismatch is defined. The fundamental basis of cybersickness still appears to be centered around the integration of sensory cues responsible for self-motion and perhaps a multisensory perspective may be able to better describe the neural mechanisms leading to sickness (Gallagher and Ferrè, 2018).

To alleviate cybersickness based on sensory mismatch, a direct solution would be to provide vestibular feedback to consolidate the conflict. One such approach has been to incorporate real movement with the VR system by allowing the observer to physically move (Chardonnet et al., 2020; Mayor et al., 2019) or by using a motion platform (Keshavarz et al., 2015; Ng et al., 2020). However, there are practical and technical limitations with motion systems and the possible repercussions of different navigation techniques in VR have yet to be thoroughly explored (Davis et al., 2014; Hildebrandt et al., 2018; Nilsson et al., 2018). Another approach is to generate an artificial sensation of movement by sending an electrical current to the mastoid process known as galvanic vestibular stimulation (GVS) (Curthoys and MacDougall, 2012; Lobel et al., 1998). Studies have shown that synchronizing the stimulation with the speed and direction of visual motion (Cevette et al., 2012) or applying the stimulation intermittently and during changes in visual motion (e.g., sharp turns and curves in a driving simulation) (Gálvez-García et al., 2015; Reed-Jones et al., 2007) can reduce experiences of sickness symptoms.

More recently, it has been observed that the vestibular stimulation does not necessarily need to be mimicking a real motion or synchronized to match the visual motion to reduce sickness. Noisy vestibular stimulation using bone-conducted vibrations to the mastoid process (Weech et al., 2018a), or using bilateral random noisy GVS (Weech et al., 2020) have both been shown to be effective at reducing sickness symptoms. This effect may be attributed to a sensory reweighting mechanism, in which the noisy vestibular stimulation leads the central nervous system (CNS) to down weight the unreliable vestibular signal and rely more on the visual system, hence reducing the apparent sensory conflict (Gallagher and Ferrè, 2018; Weech et al., 2020). It has also been reported that vection latencies are reduced in the presence of noisy vestibular stimulation (Weech and Troje, 2017) and that exposure to optic flow in VR can modulate sensitivity to vestibular input and vestibular processing (Gallagher et al., 2020; Gallagher et al., 2019). This further supports the notion of a shift in sensory weighting and the consequences for motion perception. Sensory reweighting has been predominantly observed in multisensory perception and postural control based on optimal integration around the reliability and functional significance of the sensory signals (Assländer and Peterka, 2014; Carver et al., 2006; Ernst and Banks, 2002; Ernst and Bülthoff, 2004). However, whether this mechanism has any relationship with cybersickness and sensory conflict has yet to be established. Barrett and Thornton (1968) found that observers who were more sensitive to bodily as opposed to visual cues were more likely to experience sickness in an active driving simulator, suggesting that an individuals’ “perceptual style”, a tendency to rely on one sensory modality over another, can be a possible predictor of sickness.

To assess this, however we also need to better understand how the CNS responds to VR and to determine what are some of the potential aftereffects of being exposed to VR. Most research in the field of VR primarily focuses on the mechanisms and factors responsible for inducing cybersickness and how to mitigate them, while research using psychophysical techniques has predominantly used VR as a tool to generate and present highly controlled stimuli. It is known that cybersickness symptoms can persists well after the VR exposure [see (Dużmańska et al., 2018) for a review], however the consequences of this persistence for sensory perception and behaviour have not been thoroughly explored. It has been reported that while experiencing a typical VR game (i.e., commercially available) does not affect reaction times across response activation or inhibition (Smith and Burd, 2019), it may impede learning in other cognitive tasks such as spatial orientation and the ability to visualize or mentally rotate objects, as well as sustained attention (Varmaghani et al., 2022). It has also been shown that after experiencing a VR game, decision-making in a choice reaction time is slower and is likely driven by a decrease in attention (Szpak et al., 2019). Furthermore, the accommodation reflex of the eyes which is responsible for changing focus in depth is affected by VR and is correlated with ratings of sickness (Szpak et al., 2019). Here, we aim to directly examine the effect of a typical experience of a VR game not only in regard to cybersickness, but also for sensory perception.

The subjective visual vertical (SVV) is a perceptual measure of verticality consisting of orienting a line presented visually towards the perceived direction of gravity or upright. This task has been shown to represent the integration of multisensory signals involving visual, vestibular and body cues (Dyde et al., 2006). The vestibular system signals the direction of upright through the mechanical properties of the hair cells within the otolith organs and inertia from gravity, while the orientation of the body axis also provides a reference to the direction of upright known as the idiotropic vector (Mittelstaedt, 1983). When in an upright orientation, the vestibular and body signals of vertical are aligned and the perception of upright measured using the SVV task is relatively accurate to true vertical (De Vrijer et al., 2009; Dyde et al., 2006). However, when the vestibular and body cues are dissociated by tilting the head and body from upright, the SVV estimate shifts towards the longitudinal body axis indicating a bias in the perception of upright (Alberts et al., 2016b; Aubert, 1861; Clemens et al.; 2011, De Vrijer et al., 2009; Dyde et al., 2006). It has been shown that this shift in the estimate of the perceived vertical is representative the relative changes in the contributions of the vestibular and body cues and can be modelled by a reweighting or Bayesian model (Alberts et al., 2016b, Clemens et al., 2011, De Vrijer et al., 2009, Dyde et al., 2006). There have also been reports that the SVV is sensitive to differences between gender, expert observers such as dancers and astronauts who have highly trained vestibular sensitivities and patients with vestibular deficits or loss (Alberts et al., 2015, Barnett-Cowan et al., 2010, Beck et al., 2020, Harris et al., 2017). This suggests that the SVV task can be used as a reliable measure to observe the sensory reweighting mechanism between vestibular and body signals.

The purpose of this experiment is to investigate the aftereffects of VR on sensory perception and to determine whether the occurrence of cybersickness is associated with the sensory reweighting mechanism. Using the SVV task, we plan to observe whether exposure to varying intensity levels of virtual content will lead to changes in the contributions of the vestibular and body cues in the perception of upright. Following the sensory conflict theory, if the vestibular signals need to be down weighted to consolidate the conflict induced by exposure to VR, then we expect the estimate of the SVV to shift further towards to longitudinal body axis. Furthermore, if cybersickness is indeed dependent on an individuals’ ability to dynamically reweight their sensory signals, then we should expect to be able to observe a change estimate of upright representing the change in the relative contributions of these sensory signals based on the severity of symptoms experienced. We hypothesize that there will be a larger shift in the estimate of the SVV after exposure to a high intensity VR experience compared to a lower intensity experience and we also hypothesize that this change in the perception of upright will be related to severity of cybersickness symptoms. We predict that an observer who experiences a smaller or no difference in sickness severity between the two experiences should display a larger shift in the SVV estimate compared to an observer who experiences more severe symptoms of sickness. Lastly, we aim to explore whether there are gender or other demographic differences on the perceived estimate of upright and cybersickness.

## Methods

### Participants

Data was collected from thirty-one participants (16 males and 15 females; mean age = 25.6, SD = 3.9) recruited from the University of Waterloo. Seven of the participants were volunteers, while the remaining participants were remunerated $10/h for their participation. All participants reported having no auditory or vestibular disorders and had normal or corrected-to-normal vision. This experiment was reviewed and received ethics clearance through a University of Waterloo Research Ethics Committee and all participants gave their informed written consent in accordance with the guidelines from the committee.

### Apparatus and Stimuli

The Oculus Quest system was used to provide the VR content in this experiment. The Oculus Quest has a dual OLED display with a resolution of 1600×1440 (per eye) at 72 Hz refresh rate. It is a completely wireless system featuring 6 degrees-of-freedom (6DOF) inside out tracking allowing the user to freely move around. Here we allowed participants to move within a 2.4×2.4m designated play area (Fig. 1a). Two commercially available VR games were selected for the VR content - Sports Scramble and Echo VR (Fig. 1b and c). These games were chosen to elicit different levels of sickness severity based on user reviews on the Oculus Quest store and because they share comparable gameplay aspects with both games belonging to the sports genre. Sports Scramble was chosen as the low-intensity experience and consists of three main sports (baseball, tennis, and bowling) where the user is predominantly stationary both within the game and in the physical world and mainly involves to use of arm movements to catch, throw and hit a ball in each respective sport. Echo VR was designated for the high-intensity experience and puts the user in a zero-gravity arena in which they must navigate using the provided controllers in combination with rotation of their physical body to catch and throw a disc into an opposing team’s goal.

**Fig. 1.**
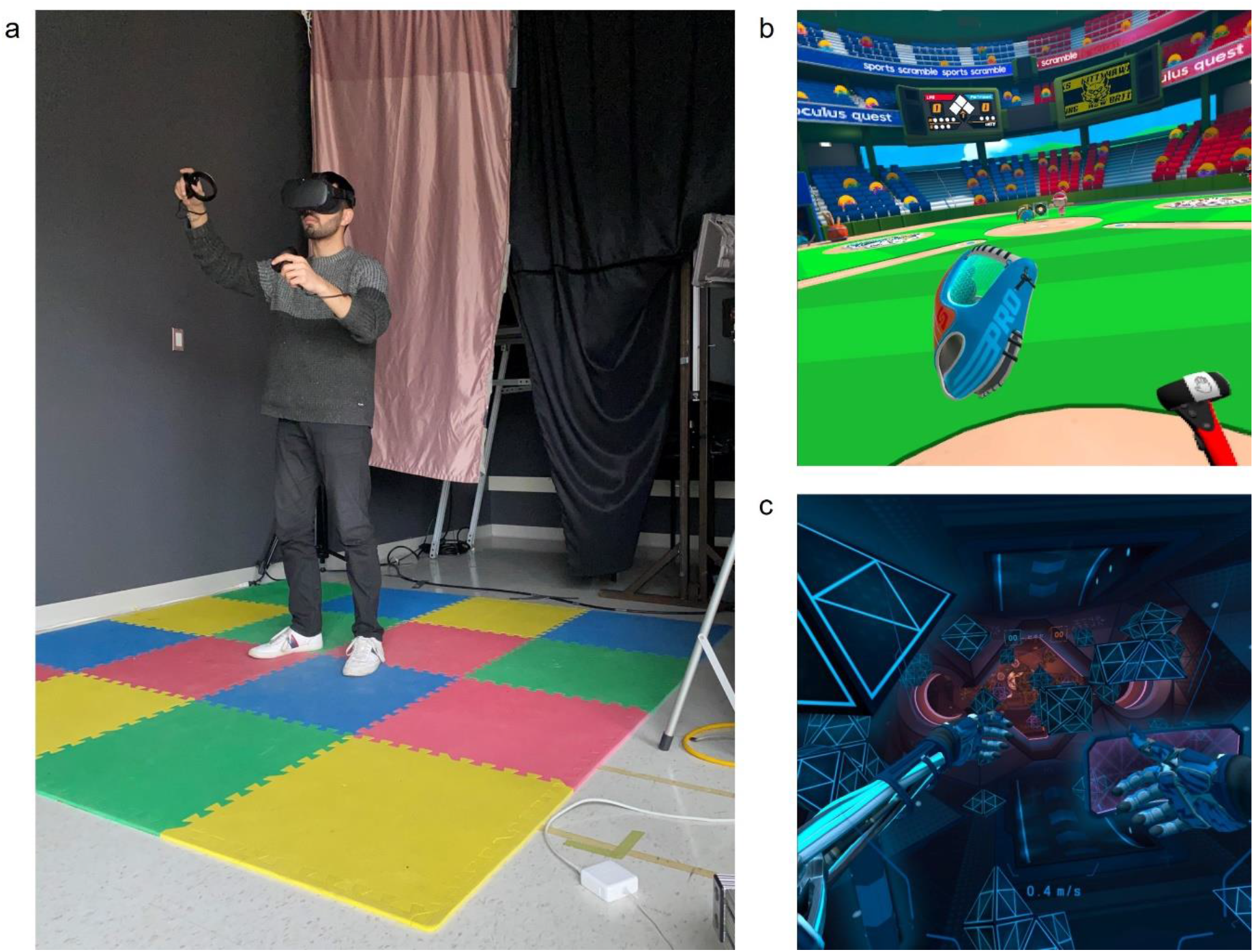
**a** Image of 2.4×2.4m designated VR play area using the Oculus Quest system. Screen capture of the game **b** Sports Scramble representing the low intensity VR and **c** Echo VR for the high intensity VR experience.

The subjective visual vertical (SVV) task was used to measure participant’s weighting of vestibular cues using the “luminous line” technique (Barnett-Cowan et al., 2010) and the method of constant stimuli. A line (3 × 0.5°) was presented for 500 ms on a neutral grey background with a central fixation point (0.45°of visual arc) of the same mean luminance on a laptop (Macbook Pro). A shroud was mounted to the display of the laptop in which participants viewed the screen through to block out peripheral vision (Fig. 2a). With this shroud, the screen subtended a 35°diameter circle at a distance of 25 cm. There were 21 line orientations from −50°to +50°in 5°increments (Fig. 2b). Each line orientation was presented seven times for a total of 147 trials, which took approximately 5 minutes to complete. Participants reported whether they perceived the line was orientated clockwise or counterclockwise relative to their perceived vertical upright using the buttons on a gamepad (Logitech Gamepad F310).

**Fig. 2.**
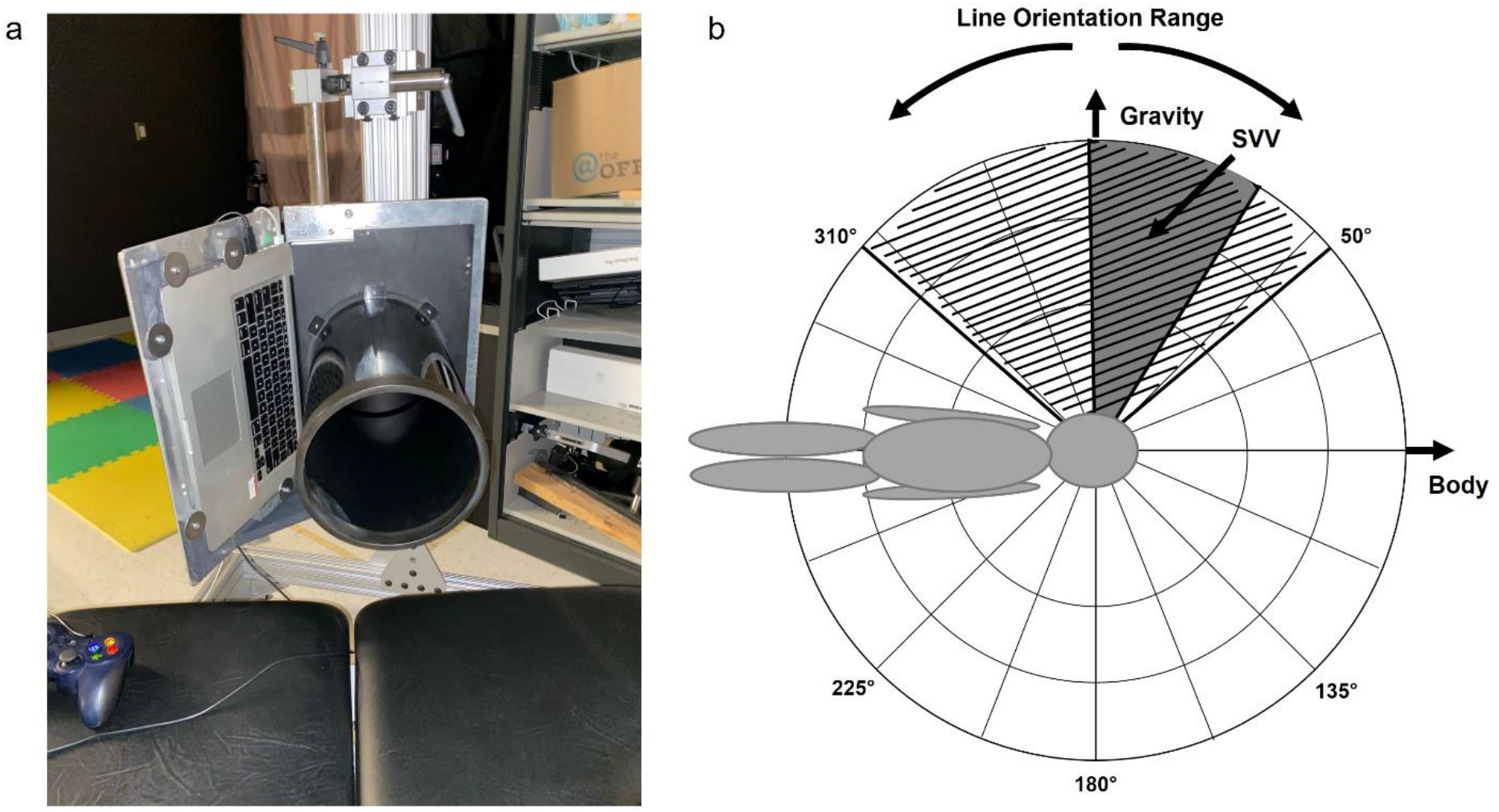
**a** Image of laptop mounted 90°clockwise on a custom stand and the circular shroud. **b** Polar illustration in earth coordinates relative to the right side down orientation. The range of the line orientations presented is highlighted by the hatched area and the shaded area indicates the range of the SVV estimates reported in the current study. The direction of upright signalled by the vestibular (gravity) and body axis cues are depicted.

### Measures

Cybersickness was assessed using the fast motion sickness scale (FMS) (Keshavarz and Hecht, 2011] to measure sickness levels immediately after each VR session was completed. The FMS is assessed by a verbal rating on a 20-point scale ranging from “0 (representing no sickness at all)” to “20 (representing severe or frank sickness)”. Participant’s gaming experience was also measured at the beginning of the experiment using a questionnaire including the following questions: “Do you have any experience with video games?”, “Do you play video games regularly?”, “On average, how many hours a week do you spend playing video games?”, “How would you describe your current level of gaming expertise?”.

### Procedures

Participants laid on their right side on an examination table with a foam pillow headrest to align their head with a circular shroud mounted on the screen of a laptop. The laptop was oriented 90°clockwise and mounted on a custom-built stand. They completed a baseline SVV task in which they had to indicate whether a series of lines presented was oriented clockwise or counterclockwise from gravity (“the direction in which a ball would fall”) via a 2-alternative forced choice task by button press on a gamepad. This took approximately 5 minutes. After completing the SVV task, the participants were guided to the VR play area where they were fitted with the Oculus Quest system. Participants played either Sports Scramble or Echo VR for 30 minutes. At the end of the VR session, participants’ sickness level was verbally assessed using the FMS scale and then participants immediately completed another SVV task for approximately 5 minutes. Participants were then given a 15-minute break and after ensuring that any sickness symptoms had subsided, they began the second VR session with the alternative game that they did not experience in the first session. The order in which participants played the two VR games were counterbalanced. After the end of the second VR session, participant’s sickness severity was measured again and followed by a final SVV assessment that again took approximately 5 minutes.

### Data Analysis

Statistical analysis was conducted using SigmaPlot 12.5 and IBM SPSS Statistics 25. A logistic function which accounted for lapses in participants’ responses (Eq. 1) was fitted to the participants’ percentage of “clockwise” responses as a function of the line orientation. Estimates of lapse rate, bias and precision were derived from the three free-parameter equation to 21 data points corresponding to the 21 line orientations tested. The halfway (50%) inflection point of the sigmoid curve (x0) represents the point of subjective equality (PSE) which here represents the SVV estimate and the standard deviation of the function (b) as the just noticeable difference (JND) (Barnett-Cowan et al., 2010). A positive SVV value indicates that the line was perceived as vertically upright when it was orientated clockwise from true vertical, while a negative SVV score indicates that the line was perceived as vertically upright when it was oriented counterclockwise from true vertical. For one participant, we were unable to fit the lapse function to their responses (*R^2^* < 0.2) and they were excluded from subsequent analyses (final sample: 15 males and 15 females; mean age = 25.5, SD = 3.9).

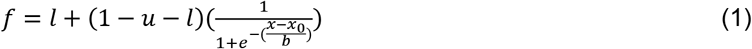

For the gaming experience survey, participants were categorized into two groups of “low” and “high” gaming experience based on their responses in the survey. Participants who had no experience with video games, did not play video games regularly or only played video games regularly for less than one hour a week were categorized into the “low” gaming experience group. Participants who played video games regularly for more than one hour a week were categorized into the “high” gaming experience group. In total, 15 participants were assigned to the “low” gaming experience group and 15 participants were assigned to the “high” gaming experience group. To assess the relationship between the SVV estimate and FMS scores, a three free-parameter Gaussian nonlinear regression model was fit to the FMS scores as a function of change in SVV from baseline after each VR exposure (Eq 2).

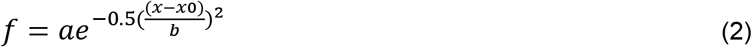

## Results

### Subjective Visual Vertical

The results showed that the SVV was shifted in a positive direction towards the body axis at all three measurement times [Baseline (M = 10.94, SE = 1.59); Low Intensity (M = 8.87, SE = 1.52); High Intensity (M = 8.07, SE = 1.68)] (Fig. 3a). To determine whether there was an effect of the VR exposure on the performance in the SVV task, a one-way repeated measures ANOVA was conducted on the SVV and JND data. Mauchly’s test of sphericity was violated for the SVV scores [*χ*^2^(2) = 7.72, *p* < 0.021] and the Greenhouse-Geisser corrected test revealed a significant main effect for the SVV scores across exposures [*F*(1.61,46.74) = 3.63, *p* = 0.043, η_p_^2^ = 0.11]. Post hoc tests with Bonferonni adjustments for multiple comparisons indicated that the SVV scores were significantly lower after the high intensity VR exposure compared to baseline (*t* = 2.61, *p* = 0.035), indicating that the SVV was set closer to gravity and farther from the body. No significant difference was found after the low intensity VR compared to baseline (*t* = 1.88, *p* = 0.129) and the high intensity VR (*t* = 0.73, *p* = 1.00) (Fig. 2a). For the JND scores, the Friedman test was performed and there was no significant main effect across exposures [*X^2^* (2) = 3.20, *p* = 0.202] [baseline (M = 4.68, SE = 0.59); low intensity (M = 4.22, SE = 0.49); high intensity (M = 5.18, SE = 0.74)] (Fig. 3b).

**Fig. 3.**
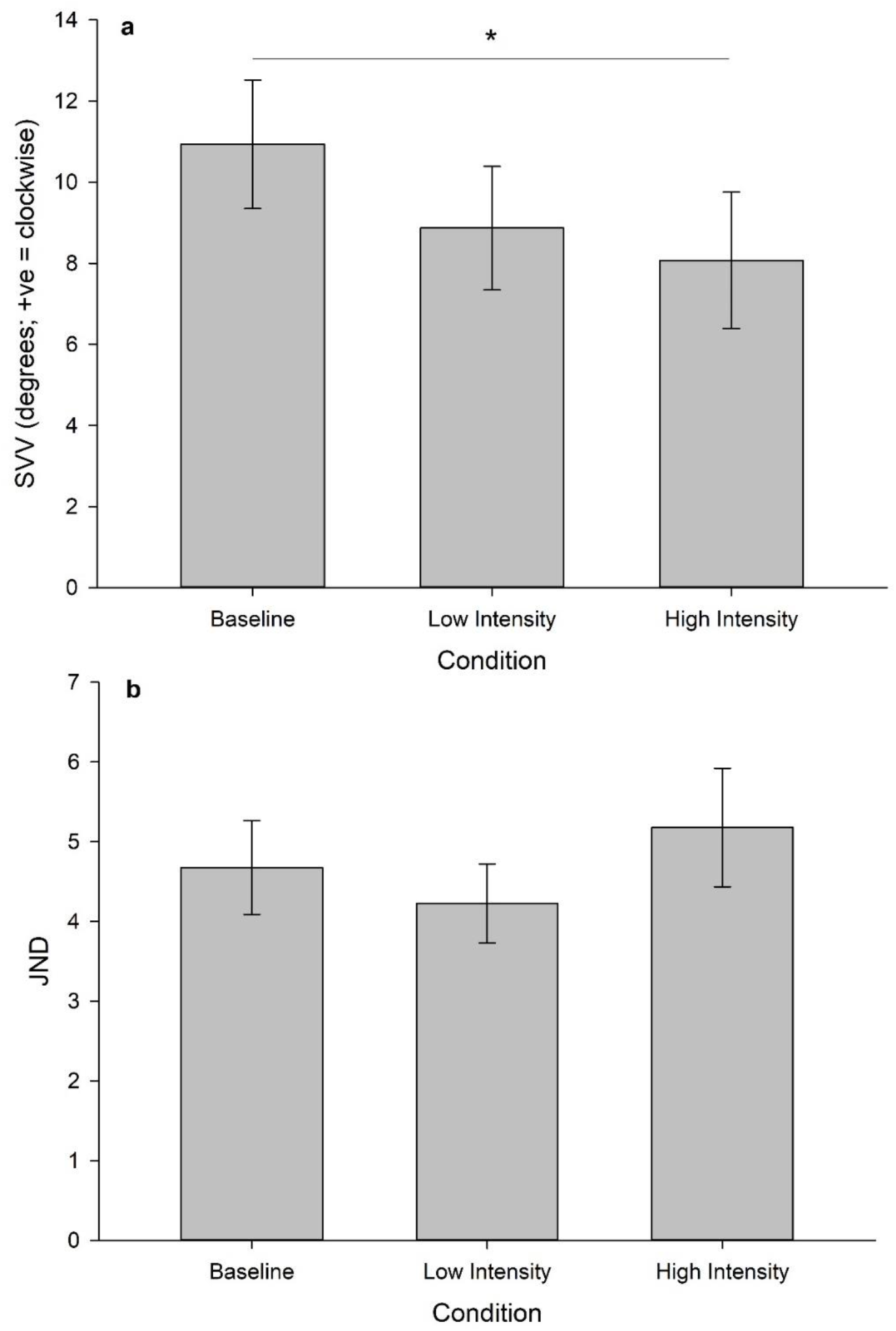
Summary of **a** subjective visual vertical (SVV) estimate and **b** just-noticeable difference (JND) between low and high intense VR exposures. Error bars are standard error.

### Cybersickness

To confirm that the low and high intensity VR experiences induced different levels of sickness severity, a Wilcoxon Signed Rank Test revealed a significant different in FMS scores (*Z* = 4.55, *p* < .001) [low intensity (M = 1.87, SE = 0.42); high intensity (M = 7.90, SE = 0.94)] (Fig. 4).

**Fig. 4.**
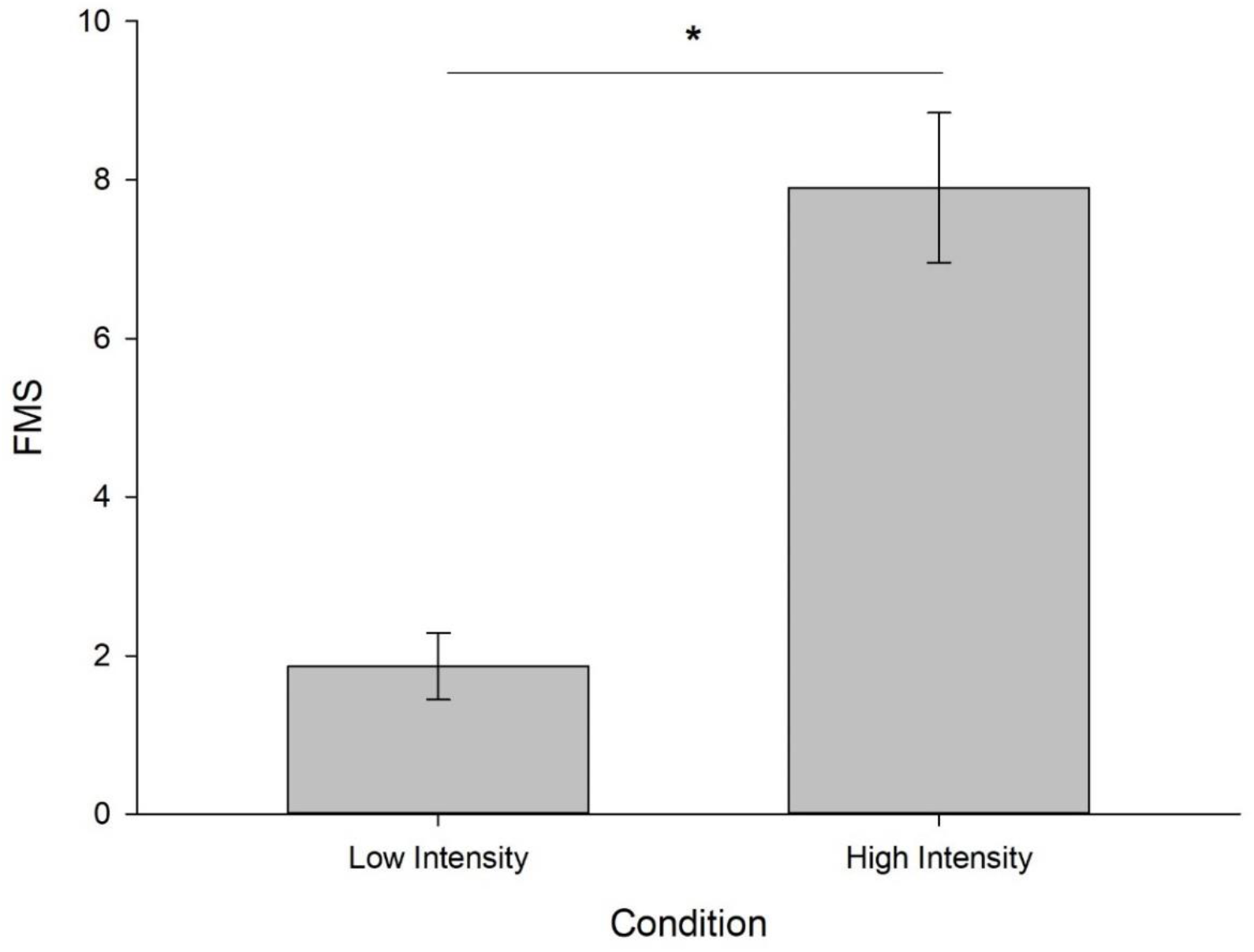
Summary of fast motion sickness (FMS) ratings between low and high intense VR exposures. Error bars are standard error.

### Subjective Visual Vertical and Cybersickness

For our main hypothesis, we wanted to examine whether a change in the perceptual weighting of gravity versus bodily cues is correlated with a change in sickness severity, represented by the SVV and FMS scores, respectively. To examine this relationship, we took the absolute values of the difference in SVV scores after the high and low intensity sessions, as well as the difference in the FMS scores (note that we did not have to take the absolute values of the FMS scores because all participants reported either the same or higher ratings after the high intensity session). As cybersickness scores are not normally distributed (Rebenitsch and Owen, 2021) confirmed by a Shaprio-Wilk test [*W*(30) = 0.90, *p* = 0.008], we conducted a Spearman’s rho correlation and found that there was no significant correlation between the absolute SVV difference and FMS difference between the low and high intensity sessions [*r* (30) = −0.198, *p* = 0.294] (Fig. 5). To further investigate the relationship between cybersickness and change in the SVV, a Gaussian regression model was fit to the FMS ratings as a function of change in SVV from baseline after each VR exposure. The results of the regression revealed that the change in SVV significantly predicted 49.5% of the variance in the FMS ratings [*R*^2^ = 0.49, *F*(5,54) = 10.58, *p* < 0.0001] (Fig. 6).

**Fig. 5.**
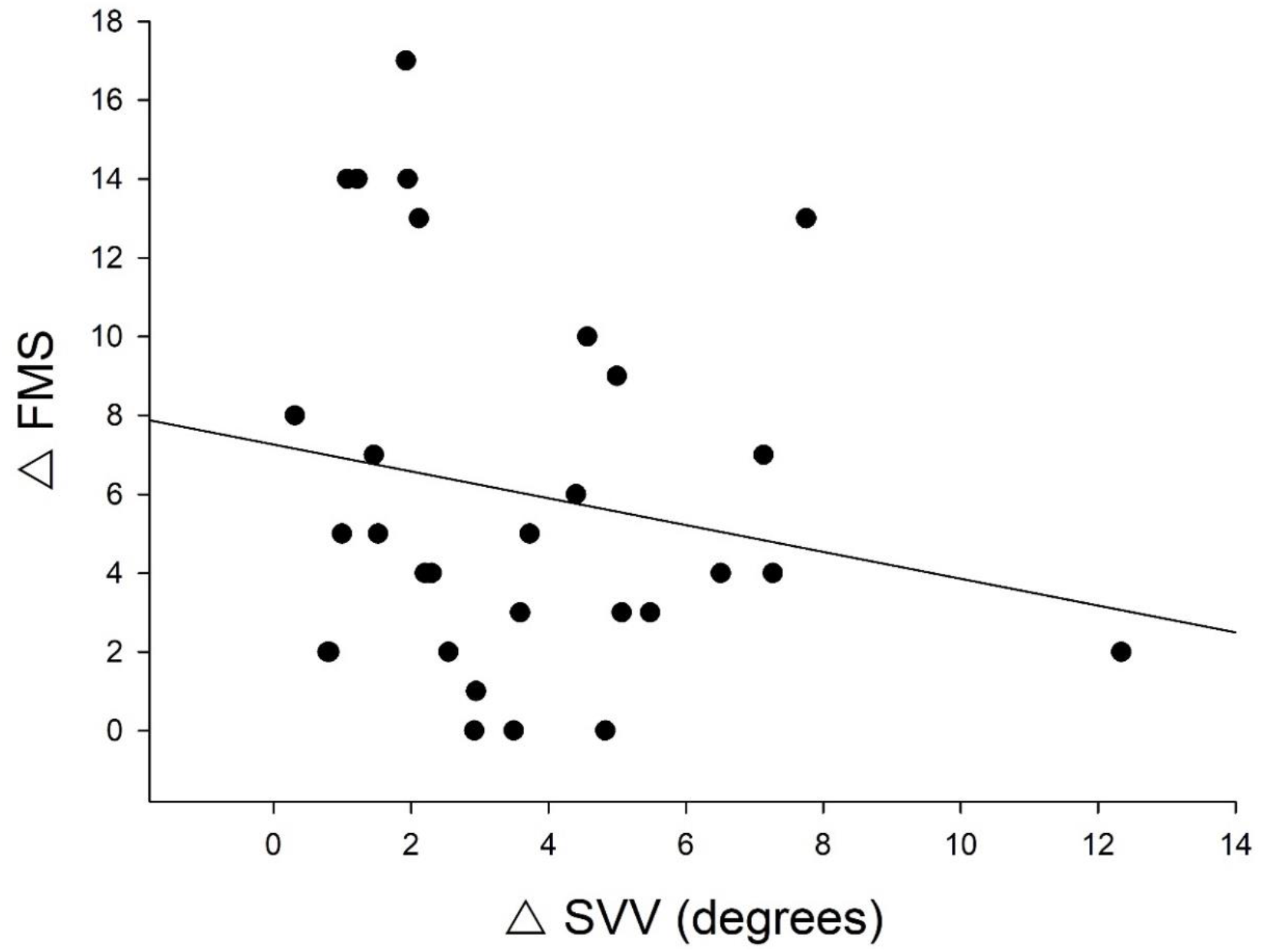
Correlation between the change in FMS score and change in absolute SVV value between the high and low intense VR exposures. The black line represents the line of best fit (*R* = −0.198).

**Fig. 6.**
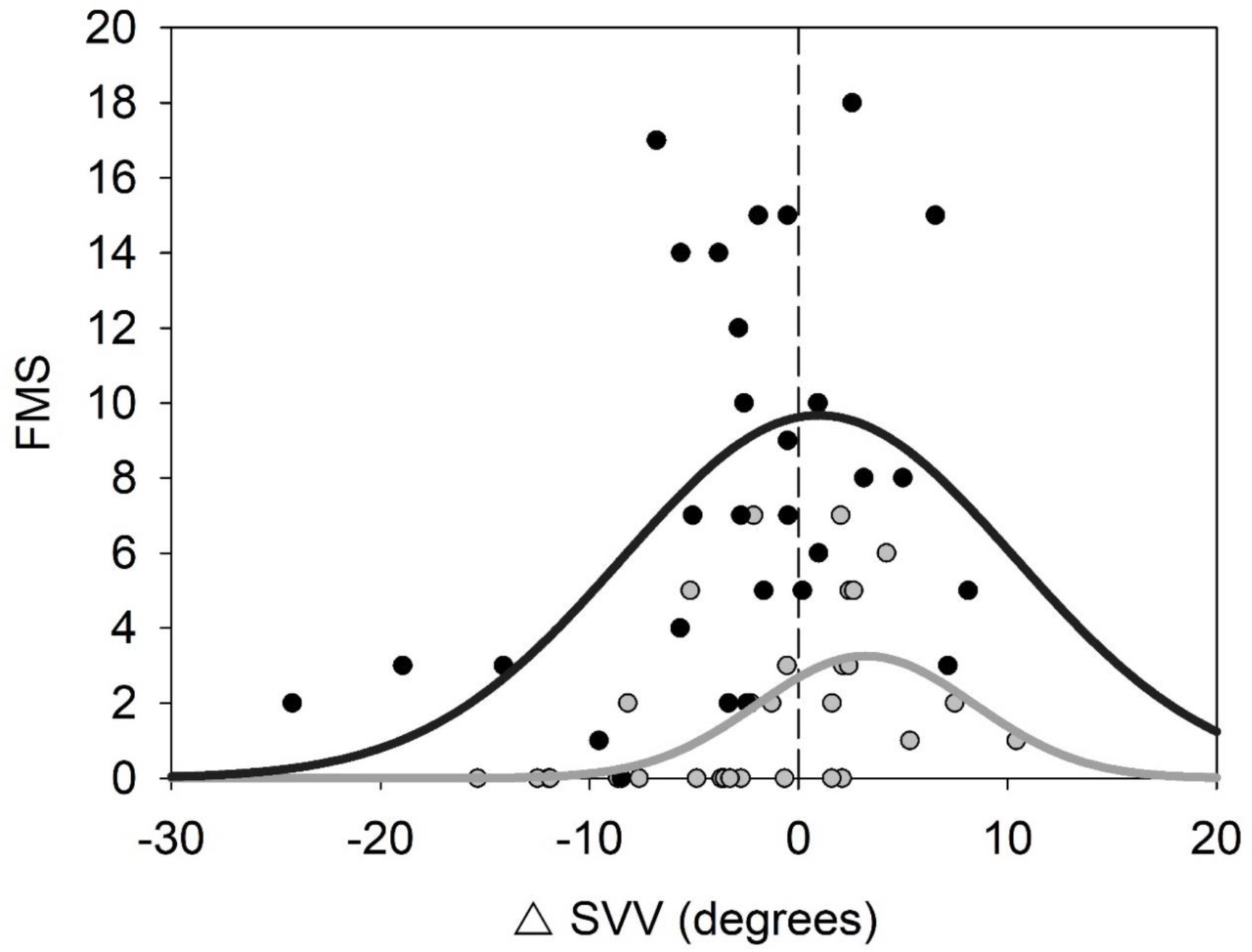
Gaussian model of FMS ratings relative to the change in SVV for both low and high intensity VR exposures from baseline. The grey-filled dots represent the data points from the low intensity VR exposure and the black-filled dots from the high intensity VR. The grey line is the Gaussian model prediction for the low intensity VR and the black line for the high intensity VR. The dashed line indicates the upper specification of the regression of no difference in the SVV from baseline.

### Individual Factors

To determine if there were any gender differences in the SVV estimates and FMS scores, the data was split between males and females (summarized in Table 1). A series of independent samples t-tests with Bonferroni-adjusted significance criteria were conducted to compare the SVV and FMS scores across the different measurement points. The results showed that there were no significant differences between males and females in the SVV [Baseline (*t*(28) = −0.195, *p* = 0.847); Low Intensity VR (*t*(28) = 0.392, *p* = 0.698); High Intensity VR (*t*(28) = −0.317, *p* = 0.754)] and FMS scores [Low Intensity VR (*U* = 89.0, *p* = 0.312); High Intensity VR (*t*(28) = −1.099, *p* = 0.281)]. The same procedure was performed to examine the effects of gaming experience between participants with low and high gaming experience (summarized in Table 1). The results revealed no significant differences in the SVV [Baseline (*t*(28) = −0.195, *p* = 0.772); Low Intensity VR (*t*(28) = −0.943, *p* = 0.354); High Intensity VR (*t*(28) = −1.005, *p* = 0.323)] and FMS scores [Low Intensity VR (*U* = 72.0, *p* = 0.079); High Intensity VR (*t*(28) = 0.383, *p* = 0.705)].

**Table 1.**
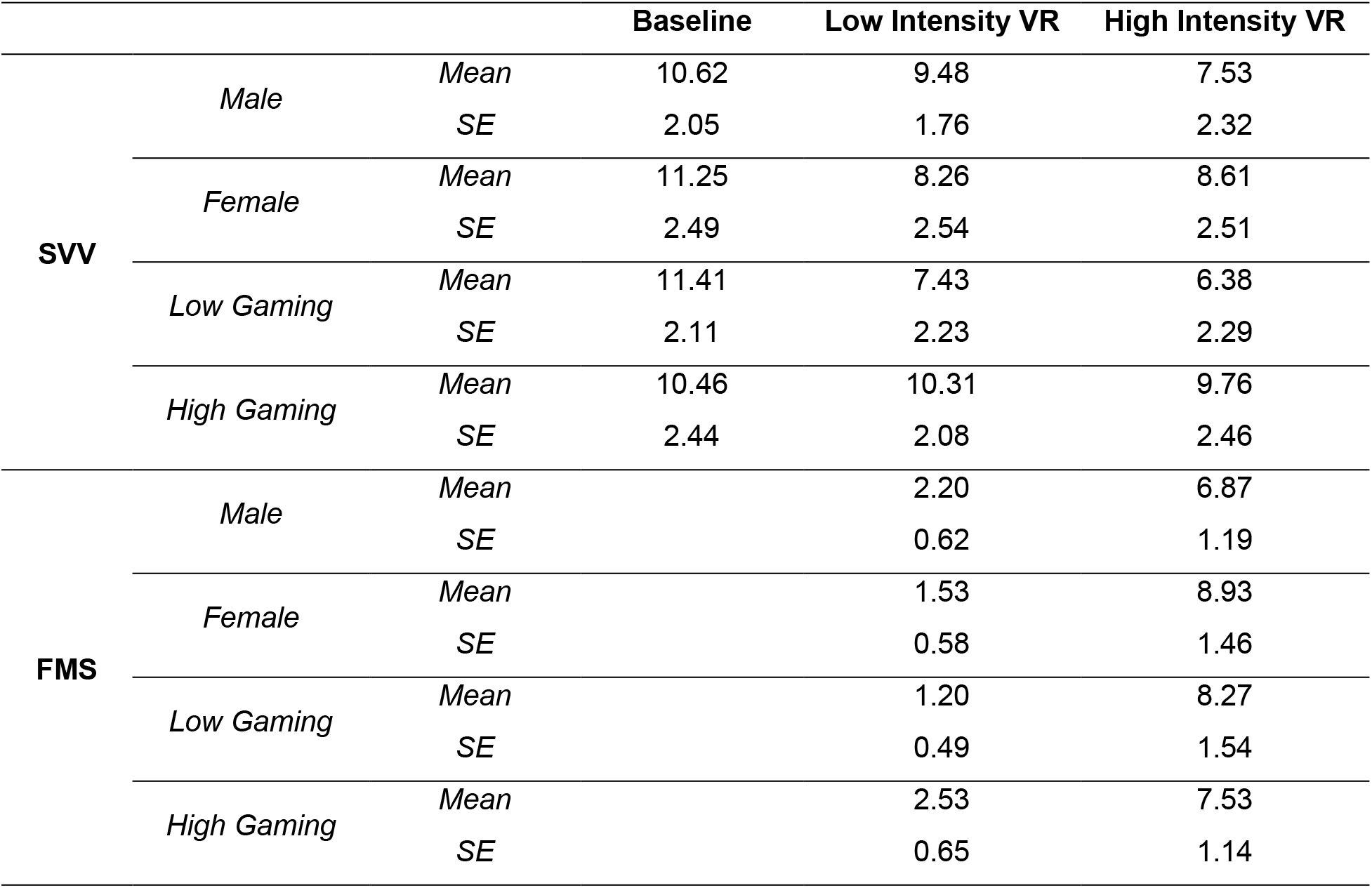
Summary SVV and FMS split by gender and gaming experience measured at baseline and after the low and high intensity VR exposures.

## Discussion

The purpose of this experiment was to investigate the effects of different VR exposures on sensory reweighting and whether there is a relationship between sensory reweighting and cybersickness severity. Sensory reweighting was observed through the systematic bias in subjective visual vertical (SVV) task towards the axis of the body when performed in a 90°clockwise roll tilt orientation (Alberts et al., 2016b; Alberts et al., 2015; Clemens et al., 2011; De Vrijer et al., 2009; Dyde et al., 2006). We found that after participants were exposed to 30 minutes of an intense VR experience there was a significant shift in the bias of the SVV away from the body axis towards the gravitational upright indicated by a significant decrease in the PSE. This finding suggests that either the contribution of the vestibular cue increased, the contribution of the idiotropic body cue (Mittelstaedt, 1983) decreased or a combination of both occurred in subjective vertical after the VR experience. Participants also reported significantly higher sickness ratings in the high intensity experience and although the correlation between the change in SVV and difference in sickness severity between the two VR experiences was not significant, there appears to be a trend for participants who experience a lower difference in ratings of sickness to have a larger shift in their estimate of the SVV compared to those who experience a greater difference in sickness.

Our results for the SVV being biased towards the body axis agrees with previous findings for the A-effect or under compensation for roll tilt on the perceived upright based on the line orientation (Alberts et al., 2016b; Alberts et al., 2015; Aubert, 1861; Clemens et al., 2011; De Vrijer et al., 2009; Dyde et al., 2006). This bias is based on the idea that body tilt introduces noise into the vestibular system (Clemens et al., 2011), presumably from the mismatch between the vestibular (otolith) cues and gravity. According to the Bayesian reweighting model, the decreased reliability of the vestibular cue leads to an increased reliance on the body cue of upright and along with the influence of a prior for each sensory modality, produces an estimate of upright based on a combination between the two signals (Clemens et al., 2011; Dakin and Rosenberg, 2018; De Vrijer et al., 2009; Dyde et al., 2006). In addition to the bias in the SVV, we also found that there was a significant shift in the SVV towards the vestibular cue after exposure to high intensity VR. Previous reports of a shift in the bias of the SVV have been found with changes in the degree of roll tilt orientation (De Vrijer et al., 2009) or bilateral vestibular loss (Alberts et al., 2015) leading to a shift in sensory reweighting, however in the present study the orientation of the observers remained consistent and only healthy participants were recruited. Another potential explanation could have been practice effects from completing the same SVV task repeatedly, however we had a subset of participants (n = 11) perform a second baseline task and found no significant difference from the initial baseline. Furthermore, the order of the two different VR experiences were counterbalanced across participants and no significant differences were observed for the SVV after the low intensity VR experience. This suggests that the significant shift in the SVV found was likely due to the exposure to the high intensity VR experience.

Following the Bayesian framework, the current results could be described as either an increase in vestibular cue weighting, a decrease in body cue weighting or a combination of both based on the change in reliability of each individual sensory signal. The current design with whole body roll tilts does not allow for us to investigate or model the contributions of each individual sensory cue in the estimate of upright. However, previous studies have found an increased reliance of the SVV on vestibular signals when dissociating the contributions of vestibular and body cues using comparisons with estimates of subjective body tilt (SBT) and head-on-body tilts. The SBT is thought to be mainly represented by body-in-space sensors but can also be derived through an indirect pathway from head-in-space coordinates modulated by the head-on-body representation and vice versa for the SVV (Clemens et al., 2011). It has been reported that the relative contributions of the sensory signals for upright differ for each of these task (Clemens et al., 2011) and while body cues through indirect pathway previously described can make up for the estimate of upright when the vestibular function is compromised, it cannot completely make up for the missing vestibular cues (Alberts et al., 2015). In another study by Alberts et al. (2016b) vestibular and somatosensory contributions to the SVV was dissociated by examining performance in the task with body tilt compared to head-only tilt. There was an increased biased with both body and head tilts, however the systematic error in SVV was found to be a function of head-on-body tilt rather than body tilt. This suggests a larger contribution of head sensors through the vestibular and otolith signals and possibly neck proprioception on the SVV over the body cues like previous reports of SVV operating in a head-fixed reference frame (Tarnutzer et al., 2010). Taken together, these findings provide some evidence to suggest that the shift in SVV observed in the current study is likely a result of an increase in vestibular weighting rather than a decrease in reliance on body cues.

Why is the SVV shifted after exposure to VR and more specifically why is the weighting of vestibular cues increased? Harris et al. (2017) measured the SVV and perceptual upright in astronauts pre-flight and after returning from long term exposure to microgravity in space. There were no significant differences between the astronauts and a control group for the shift in SVV pre- and post-flight, however variance in the SVV without visual orientation cues were significantly increased in astronauts post-flight. While we cannot directly compare the current results with these findings, this study showed that the SVV is sensitive to the balance between vestibular and body cues in the representation of upright and can reflect the changes in sensory reweighting between these sensory signals. It is plausible then that the shift in SVV found in the present study stems from potential sensory reweighting that occurred when observers were exposed to VR. It has been shown that VR can modulate vestibular sensitivity and processing (Gallagher et al., 2020; Gallagher and Ferrè, 2018) and our results agree with these findings that the reliance on vestibular signals were altered after exposure to VR. As to why it seems that the weighting of vestibular cues increased, we know that sensory conflict between vestibular and visual cues in VR is a potential cause of cybersickness (Cobb et al., 1999; Howarth and Costello, 1997; Keshavarz et al., 2015; Kim et al., 2020) and minimizing or consolidating this mismatch can alleviate symptoms (Ng et al., 2020; Teixeira and Palmisano, 2021; Weech et al., 2018a; Weech et al., 2020).

While previous strategies have focused on down-weighting the vestibular signal to rely more on the visual motion signals present in VR, it is also possible that the vestibular signals could be up-weighted to promote physical stability. According to the postural instability theory in cybersickness, sickness is correlated with postural instability or when people are naturally unstable or unable to maintain control of their posture when exposed to VR (Arcioni et al., 2019; Assländer and Peterka, 2014; da Silva Marinho et al., 2022; Rebenitsch and Quinby, 2019; Risi and Palmisano, 2019). While we did not observe a significant correlation between the difference in sickness rating and difference in SVV, there was a trend towards lower difference in sickness with greater changes in SVV and sickness severity was significantly greater in the high intensity VR experience. Thus, the weighting of vestibular cues may have increased to promote postural stability modulated by cybersickness from the VR exposure, however future work including a measure of posture control during the VR experience is needed to explore this hypothesis.

To further investigate the relationship between cybersickness and the SVV, a regression model was used to explore whether cybersickness ratings could be predicted by the change in performance in the perceptual task. A recent review modelling the severity and incidence of cybersickness in the literature found that demographic information can explain 44.2% and hardware and software factors can explain up to 55% of the variance in cybersickness (Rebenitsch and Owen, 2021). Previous research has also attempted to predict cybersickness using a similar regression approach with various factors. Kim et al. (2005) used a stepwise regression model and a battery of physiological measures to determine that a combination of autonomous variable (heart period), electroencephalogram (EEG) activity representative of an increase in cognitive demand and stress response and past motion sickness susceptibility was able to predict 46% of the variance in the severity of cybersickness symptoms measured using the Simulator Sickness Questionnaire (SSQ). A similar report using the same modelling approach and a different range of physiological measures found that stomach activity, blinking and breathing rates can explain 37.4% of the variance in cybersickness measured by the SSQ (Dennison et al., 2016). In addition to physiological factors, it has also been shown that vection strength (Nooij et al., 2017) and eye movement behaviours (Chang et al., 2021) can be strong predictors of sickness (48% and 34.8% respectively).

However, in all these studies the participants were limited in their ability to interact with the virtual environment due to restrictions from the measurement devices or just passively viewing a virtual environment without user control. In a study more comparable to the current design by Weech et al. (2018b), balance control, vection and vestibular measures were taken before completing two different VR tasks of varying intensities and found that their model predicted 37% of the variability in cybersickness measured by the SSQ with a strong contribution (16%) from postural stability. In the present results, 49.5% of the variance in cybersickness measured using the FMS scale was predicted by the change in SVV performance alone. It should be noted that previous models used the SSQ as the primary sickness measures which calculates a weighted score based across multiple symptoms, while here the FMS score on a 21-point Likert rating scale was used. This indicates that the interaction between visual, vestibular and body cues in the perception of upright is a strong predictor of cybersickness and suggests that sensory reweighting may be an important underlying mechanism.

Both the SVV and cybersickness are subjective measures that can greatly vary between individuals as indicated by the magnitude of the variance observed in the results. As previously mentioned, a model of existing data in the literature showed that demographic information can explain 44.2% of the variance in cybersickness (Rebenitsch and Owen, 2021). For these reasons, it is important to consider demographic and individual factors when investigating cybersickness to try to account for these potential individual differences. For the SVV, gender differences have been previously reported in which females are more likely to be influenced by the orientation of a visual background when upright [Barnett-Cowan et al., 2010] and mixed results have been found with expert observers, where dancers were found to be more susceptible to the vestibular noise generated by head tilts (Beck et al., 2020) and astronauts performed no different than a control group on earth (Harris et al., 2017). Recent reviews have also highlighted various factors related to individual, hardware, content, and experimental paradigm differences leading to the vast differences observed for susceptibility to cybersickness in the literature (Howard and Van Zandt, 2021; Tian et al., 2022).

There have been contrasting reports regarding gender differences with females being more susceptible to cybersickness, but recent findings have begun to suggest that this may have been due to improper fitting of the HMDs to the inter-pupillary distance (IPD) of females, as well as differences in sensitivity to motion parallax and 3D visual acuity (Allen et al., 2016; Fulvio et al., 2021; Stanney et al., 2020). Another interesting factor is gaming experience, in which it appears that people with more gaming experience may be less susceptible and recover faster from sickness (Curry et al., 2020; da Silva Marinho et al., 2022). The genre or content of video games can also influence the occurrence of cybersickness with action and first-person shooter games being more likely to induce sickness than adventure games, possibly due to the greater amount of fast angular motions in those genres (da Silva Marinho et al., 2022; Rebenitsch and Owen, 2021). This may be related to video game experience, as users with more experience may have reduced susceptibility to cybersickness due habituation or familiarity with the characteristics of the game content or improved visuospatial skills (Smyth et al., 2021). In the present study, we did not find any significant gender differences in SVV or cybersickness as we used a more recent HMD model with IPD adjustments (Oculus Quest) and anecdotally, no participants reported adjustments issues with the lens. We also did not find a significant effect of gaming experience, although it is interesting to note that most of the participants that reported “low” gaming experience were female (11 out of 15) and a majority of the “high” gaming experience were male (11 out of 15). Future work with a larger and balanced sample size is needed to properly examine these demographic factors.

Sensory conflict in VR primarily involves the visual, vestibular, and proprioceptive systems, however in the present SVV paradigm, we did not include a measure for the contributions of vision. Visual feedback has been shown to have a significant influence on the perception of upright, with visual orientation cues in the form of a static scene or frame background when vestibular and body cues are not aligned (Alberts et al., 2016a; Alberts et al., 2019; Alberts et al., 2018; Barnett-Cowan et al., 2010; Dyde et al., 2006; Harris et al., 2017). Since we did not provide any visual orientation cues, our interpretation of the current results is limited to the vestibular and body axis cues. Having the contributions of vision would have allowed us to model the reweighting of the sensory signals involved in the perception of upright (Alberts et al., 2016a; Dyde et al., 2006) and provide a more comprehensive understanding of what is occurring to the sensory systems after VR exposure. Another limitation of the current study is potential carry over effects between the VR exposures given the time that participants had to rest and recover from any sickness symptoms in between. We chose a 15-minute break following the guidelines of the Meta Quest Health & Safety Manual, however the persistence of sickness symptoms is not fully established and has been reported to last from anywhere between 10 minutes to up to 4 hours (Dużmańska et al., 2018; Szpak et al., 2020). A multitude of factors can contribute to the persistence of the symptoms such as the individual’s susceptibility, the state of the user, exposure time, the VR content, the hardware used and along with relying on subjective reports on symptoms makes it extremely difficult to control for each participant. One solution could have been to have participants conduct the second VR exposure and associated SVV task on a different day, however this would prevent the comparison of the SVV between the two conditions. Lastly, the VR content used in the current study were commercially developed games and how each participant interacted with the game content could only be controlled to a certain extent across participants. This limits the depth of our interpretation of the findings regarding the specific aspects of the VR exposure beyond the sickness ratings that led to the observed results.

## Conclusion

In the current study, the relative contributions of vestibular and body cues in the perception of upright were examined after exposure to two VR experiences of different intensity levels. Furthermore, we investigated whether the perception of upright and susceptibility to cybersickness were related based on a common sensory reweighting mechanism. The results showed that exposure to a high intensity VR experience resulted in a larger shift in the perception of upright towards the vestibular signal compared to a low intensity VR content. This finding suggests that the sensory weighting of the vestibular cue in perceiving upright increased after the high intensity VR exposure. However, there was no significant correlation between the change in perception of upright and susceptibility to cybersickness quantified as the change in sickness ratings between the two VR experiences. Despite this result, there still appeared to be a trend towards the hypothesis that there is a shared sensory reweighting mechanism for some participants and the change in performance in the perceptual task between the VR exposures was able to predict 49.5% of the variability in the sickness measure. Future research can try to better capture this relationship and the overall findings from this study provide important implications for potential aftereffects of general use of VR for entertainment in a more typical setting.

